# Adaptive convergent evolution of genome proofreading in SARS-CoV2: insights into the Eigen’s paradox

**DOI:** 10.1101/2021.09.11.459886

**Authors:** Keerthic Aswin, Srinivasan Ramachandran, Vivek T Natarajan

## Abstract

Evolutionary history of coronaviruses holds the key to understand mutational behavior and prepare for possible future outbreaks. By performing comparative genome analysis of nidovirales that contain the family of coronaviruses, we traced the origin of proofreading, surprisingly to the eukaryotic antiviral component ZNFX1. This common recent ancestor contributes two zinc finger (ZnF) motifs that are unique to viral exonuclease, segregating them from DNA proof-readers. Phylogenetic analyses indicate that following acquisition, genomes of coronaviruses retained and further fine-tuned proofreading exonuclease, whereas related families harbor substitution of key residues in ZnF1 motif concomitant to a reduction in their genome sizes. Structural modelling followed by simulation suggests the role of ZnF in RNA binding. Key ZnF residues strongly coevolve with replicase, and the helicase involved in duplex RNA unwinding. Hence, fidelity of replication in coronaviruses is a result of convergent evolution, that enables maintenance of genome stability akin to cellular proofreading systems.

## Introduction

SARS-CoV-2, the causative pathogen of CoVID-19 disease belongs to coronaviridae family of nidovirales order. CoVID-19 has infected approximately 190 million people worldwide leading to around 4 million deaths and still counting(int 2020). Coronaviruses feature among the top three family of viruses with zoonotic spillover potential **(Sup Fig 1)**(Grange et al. 2021). Members of this family have caused major outbreaks of respiratory syndromes in recent decades and are the cause of trepidation for the ongoing pandemic(NIAID;NIH). The origin of this virus is highly debated and appears to have arisen from the bat coronaviruses. The mutational events that would have led to the breach in species specificity appear to lie with the receptor binding region of the encoded spike protein. This as well as additional changes necessary for the virus to adapt to human host needs chance mutational events primarily arrived during the replication of its genome. In recent times RNA editing as well as recombination have emerged as additional source of variability. However, the primary source of mutation still lies with the RNA-dependent RNA polymerase. Emergence of SARS-CoV2 mutants, many of which challenge the protection offered by vaccines, necessitates understanding the evolutionary history of coronaviruses in the context of mutagenesis of their RNA genome.

The error rate associated with self-replicating molecules poses a upper limit suggested by Eigen’s paradox on replication fidelity and genome size threshold(Nga et al. 2011; Smith et al. 2014). Most RNA viruses obey this and are relatively small in size. However, nidoviruses have emerged as an exemption and many of the coronaviruses have a large genome size of the order of around 25 to 30 kb. Unusually large genome size in coronaviruses is attributed to replication fidelity conferred by proof-reading exonuclease (ExoN)(Minskaia et al. 2006; Lauber et al. 2013; Ogando et al. 2019). This enzyme functions alongside RNA replicase in a manner similar to DNA proofreading systems and has emerged as a solution for the paradox. However, its origin and molecular functioning has remained enigmatic.

In this study we perform comparative genome analysis and identify motifs in ExoN that are crucial to RNA proofreading and help sustain a large SARS-CoV2 genome. Structural comparisons and simulation studies ascribe a molecular functionality to this RNA proofreading enzyme which suggests distinct and more intricate proofreading compared to its DNA counterpart. Thereby, genome guided structural studies pave the way to understand the RNA proofreading mechanism that is central to the origin of mutants of SARS-CoV2 pathogen.

## Results

### Tracing the evolution of human RNA pathogens by the comparative genomic analysis of SARS-CoV2

To map the genome organization, we performed comparative analysis of RNA viruses that infect humans. Predictably, surface proteins were highly divergent. Hence, the main focus of comparison were the non-structural proteins (NSP) that are responsible for virus survival and replication of their RNA genome inside the host cell. In **Fig 1a** the heat map represents percent similarity of NSPs compared with SARS-CoV2 and the genome size for each virus is depicted alongside. Two key observations emerged from this comparison. Enzymes involved in polyprotein cleavage and primary replication machinery were fairly well conserved and coronaviruses had significantly larger genomes compared to other viral RNA pathogens. The most conspicuous was the conservation of primary RNA dependent RNA polymerase (RdRP) encoded by NSP12 in SARS-Cov2, that was identifiable across almost all viruses barring retroviruses.

**Figure 1:**
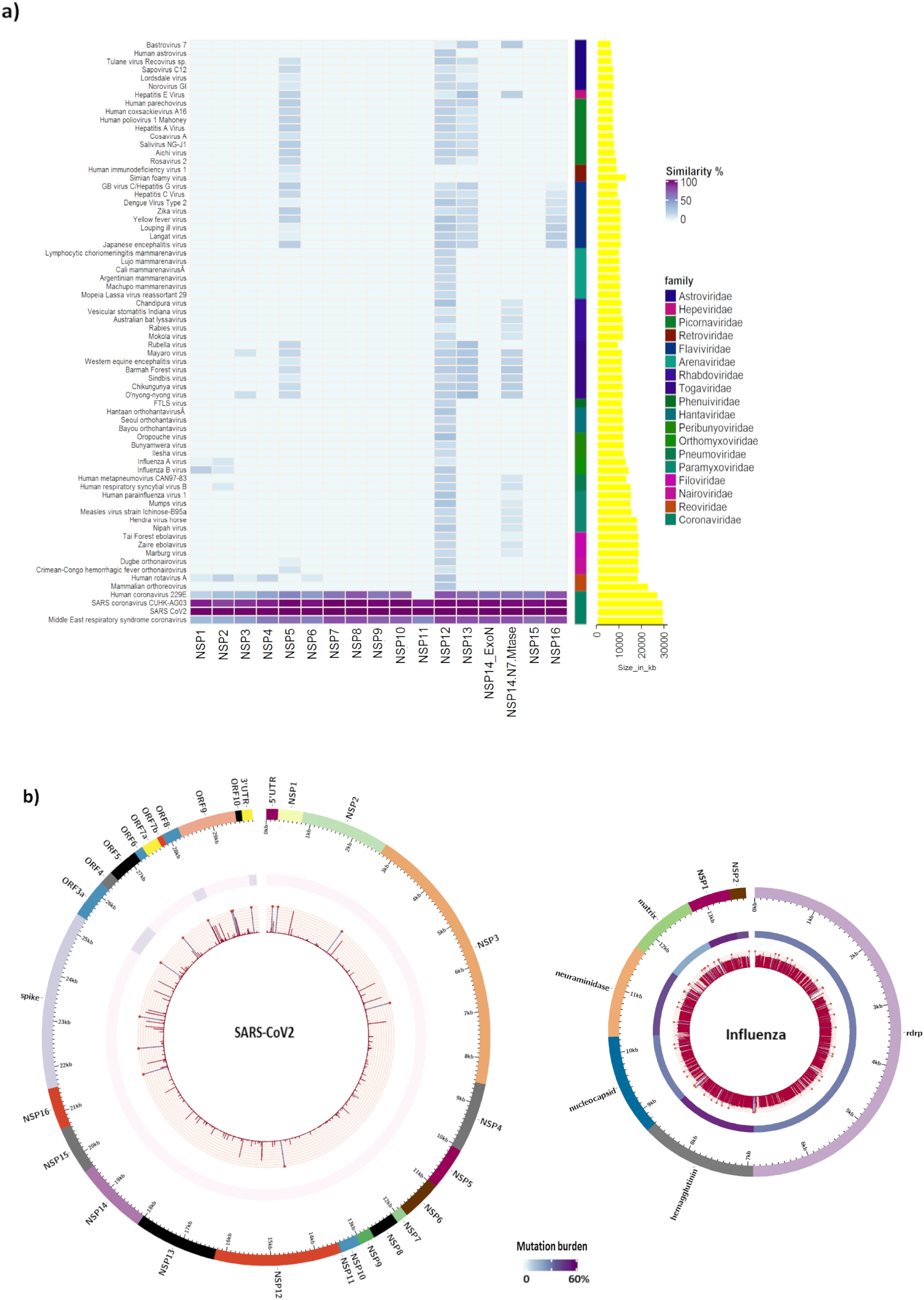
Tracing the evolution of human RNA pathogens by the comparative genomic analysis of SARS-CoV2. **a) Comparative genome analysis of Non-Structural Proteins among the known human RNA viruses** Heatmap depicts the percent similarity between SARS-CoV2 NSPs and corresponding proteins from other RNA viruses infecting humans. The respective genome size of each virus is plotted as yellow bar plots on the right. The scale bar for the heat map is depicted. b) **Mutation burden in SARS-CoV2 and H1N1 influenza** Circos plot represents mutations across the SARS-CoV2 and H1N1 influenza genomes pertaining to the current pandemic and the 2009 – 2010 pandemic respectively. The outer whorl is the genome of the virus color coded based on the encoded protein. The second whorl represents a heatmap depicting the overall mutation burden at each gene, calculated as the number of base positions with a mutation frequency more than 0.01 normalized to the gene size. The inner whorl consists of histogram representing SNP frequencies across the genome. Violet color spike with a * indicates the frequencies ≥ 0.1. Dark Magenta color spikes indicate the frequency < 0.1 The scale bar for the heat map for the second whorl is depicted.

RNA replication by the core replicase, RdRP is of low fidelity compared to DNA polymerases. Consequently RNA viruses tends to accumulate mutations at a rapid rate. A quick comparison of the mutation burden of two respiratory RNA viruses, SARS-CoV2 in the current pandemic (data from December 2019 to February 2021) with the H1N1 (2009 – 2010 pandemic) was carried out. The outcome of this comparison would be a combined reflection of the mutational capability of the virus and its selection within the population(s) by counteracting immune forces, population dynamics and advancements in the sequencing technology. The SNP frequency across the genome and mutation burden at each gene is visualized as a circos plot (**Fig 1b**). Despite being double the size and confounders against SARS-CoV2, it is evident that the smaller H1N1 influenza virus genome accumulated significantly more mutations. The burden of mutation at each gene in case of SARS-CoV2 was less than 10% whereas H1N1 influenza virus had a mutation load more than 50%. Interestingly, the structural proteins had higher mutation burden compared to the non-structural proteins, conceivably as a consequence of selection from the immune system and could explain immune-evasion. Since both these viruses replicate using the conserved RdRP, evolution of proof-reading mechanism of SARS-CoV2 appears to be built upon this core replication machinery with evolved modules that recognize mismatch and evoke exonucleolytic excision resulting in low mutation burden observed.

### Tracing the evolutionary origin of ExoN

In an earlier systematic report, by studying genomes of nidovirales order that contain coronavirus family, authors reinforced the role of proof-reading activity of ExoN encoded by NSP14 in SARS-CoV2, in contributing to larger genome size maintenance and its complexity(Chen et al. 2007; Lauber et al. 2013). However, the binary presence or absence of the proof-reading ExoN does not explain wide distribution of genome size. Especially the medium sized genomes of mesoniviridae and medioniviridae remain unexplained (**Sup Fig 2**). We traced the evolutionary origin of ExoN using NSP12 based phylogeny that revealed segregation of two major clades separating arteriviridae and tobaniviridae from other families (**Fig 2a**). Arteriviridae encompassed viruses with a consistently small genome size and lacked NSP14, whereas both tobaniviridae and coronaviridae encode ExoN and have large genomes. Based on this phylogeny, tobaniviridae appears to be the earliest ancestor to acquire a functional proofreading exonuclease. Coronaviridae ExoN encodes an associated methyl transferase domain, whereas tobaniviridae lacked this fusion in its genome, further substantiating the acquisition in this family of viruses.

**Figure 2:**
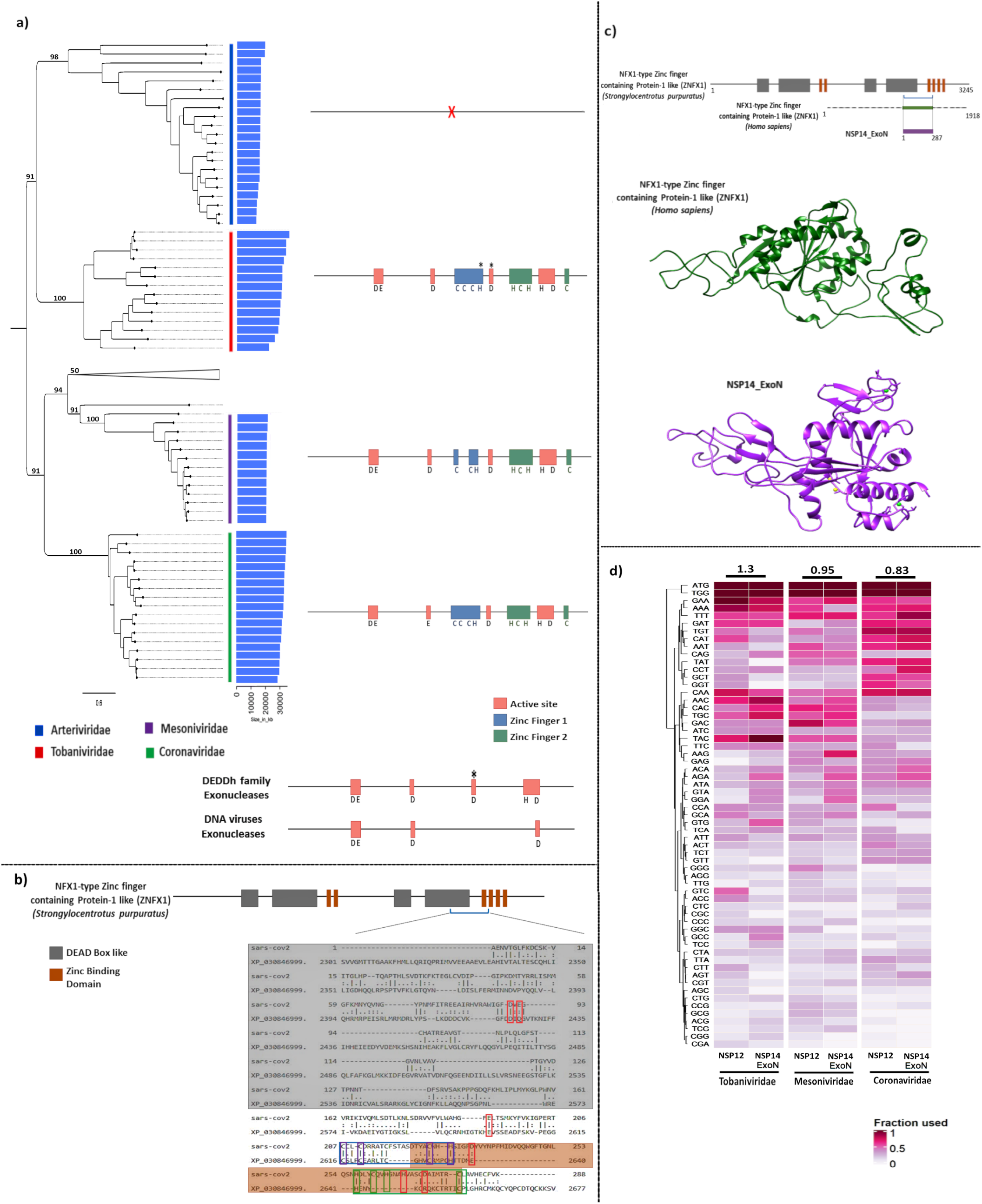
Tracing the evolutionary origin of ExoN. **a) NSP12 based phylogeny of viruses in Nidovirales order** Phylogenetic tree of Nidovirales genomes based on NSP12 is depicted. Each node represents the NSP12 from individual viruses. The corresponding families to which the viruses belong are color coded and represented. The genome size for each virus is plotted as blue bar plots. The domain architecture and key residues in ExoN across four families, Arteriviridae, Tobaniviridae, Mesoniviridae and Coronaviridae that have higher representation are depicted (to the right side). The red boxes represent the catalytic sites, blue boxes represent zinc finger 1 (ZnF1) residues and green boxes represent zinc finger 2 (ZnF2) residues. The * represents that the residue is not conserved among all the sequences analyzed. The domain architecture of DEDDh family exonucleases and DNA viruses compared to ExoN are depicted (not to scale). **b) Mapping ExoN to the common recent ancestor ZNFX1** Domain architecture of ZNFX1 and mapping of ExoN sequence is carried out to identify key residues derived from this common recent ancestor. The red boxes represent the catalytic residues, blue boxes represent zinc finger 1 (ZnF1) residues and green boxes represent zinc finger 2 (ZnF2) residues. The sequence shaded in grey is annotated DEAD box like helicase and brown region annotated as Zinc binding domain. **c) Topology comparison between ExoN and predicted structure of human ZNFX1** Human ZNFX1 (in green) and ExoN (in purple) mapping to the domains of *Strongylocentrotus purpuratus* ZNFX1 is depicted (on top). The aligned region from the predicted structure of human ZNFX1 and the ExoN backbone are depicted (at bottom). **d) Adaptation of ExoN across nidovirales traced through the codon usage pattern** The fraction of each codon used by ExoN and NSP12 for comparison, among tobaniviridae, mesoniviridae and coronaviridae are depicted as a heatmap. The Euclidean distance between NSP12 and ExoN codon usage pattern for each family is represented at the top of the heatmap.

ExoN belongs to DEDDh family of exonucleases present across all kingdoms of life including DNA viruses, and constitute enzymes involved in DNA proof-reading as well as RNA metabolism(Zuo and Deutscher 2001; Cruz-González et al. 2021). Multiple sequence alignment of ExoN with diverse homologs suggested conservation of core catalytic motif involved in exonuclease activity(Robson et al. 2020). We observed two zinc finger (ZnF) like motifs within this enzyme that are unique to nidovirales, and the crystal structure of SARS-CoV ExoN confirms the co-ordination of zinc by these two motifs(Ma et al. 2015; Lin et al. 2021). Interestingly, two of the core catalytic residues fall within the second zinc finger motif (ZnF2) (**sup Fig 3a & 3b**). Hence ExoN is structurally and possibly functionally distinct from other DEDDh enzymes, whose origin in RNA viruses is enigmatic. To identify the common recent ancestor of ExoN, position specific iterative-blast was carried out using ExoN from members of tobaniviridae. Convergence of iteration led us to a cluster of potential hits from cellular organisms. Interestingly, eukaryotic antiviral protein NFX1 type zinc finger containing protein (ZNFX1) emerged as the candidate with substantial similarity to viral ExoN (**Sup Fig 3c**).

Pattern matching had the highest score for the invertebrate homolog from purple sea urchin *Strongylocentrotus purpuratus* followed by vertebrate and fungal homologs. Pairwise alignment of SARS-CoV2 ExoN with ZNFX1 identified conservation of the two ZnF motifs and three out of six key catalytic residues. Strikingly, two of these that find orthology with ZNFX1 map to the ZnF2 domain unique to this family of exonucleases (**Fig 2b**). This common recent ancestral protein contains two DEAD like domain and four ZnF motifs and functions as a helicase with RdRP in invertebrates and fungi to cause RNA interference(Ishidate et al. 2018). The vertebrate homolog functions independent of RdRP as a dsRNA sensor and elicits an antiviral response through its core helicase domain(Wang et al. 2019). Predicted structural fold of human ZNFX1 broadly matches the structure of ExoN (**Fig 2c**). Based on these findings we propose that this common recent ancestor seems to have been acquired by nidovirales and fine-tuned to function as a proof-reading enzyme. The pattern of codon usage between NSP14 ExoN and NSP12 in tobaniviridae is dissimilar but is comparable in coronaviridae, reiterating possible ancestral acquisition followed by adaptation in this lineage (**Fig 2d**). Thereby, we speculate eukaryotic origin of this RNA virus enzyme that could have further evolved in coronaviruses to function with the core NSP12 replicase as a proofreading exonuclease. It is interesting that these viruses could have acquired an antiviral protein and mold it to strengthen the stability of their genomes.

### Identification of ZnFs role in RNA binding

Mutagenesis studies of SARS-CoV ExoN showed the importance of ZnF 1 and 2 in structural stability and exonuclease activity respectively(Ma et al. 2015; Ogando et al. 2020). The molecular function of ZnFs however remains to be explored and we hypothesized that the two ZnFs may bind RNA and help in mismatch recognition for repair. To identify the binding specificity of these two ZnFs, phylogenetic tree was constructed along with biochemically and structurally characterized zinc finger proteins that bind either single or double stranded RNA in a selective or non-selective mode of interaction(De Guzman et al. 1998; Searles et al. 2000; Méndez-Vidal et al. 2002; Hudson et al. 2004; Möller et al. 2005; Burge et al. 2014). ZnF1 clustered along with protein that binds single stranded RNA, whereas ZnF2 clustered with a protein that binds double stranded RNA (**Sup Fig 4**). Docking followed by Molecular Dynamics (MD) simulations were performed to validate the RNA binding to ZnFs. Binding pocket prediction in the modelled SARS-CoV2 NSP14 ExoN identified ZnF1 and catalytic site to be probable sites for RNA to dock (**Sup Fig 5**). The dynamics of RNA interaction with ExoN at different time points during the 300 ns simulation is represented as snapshots (**Fig 3a & 3c and Sup Fig 6a & 6c**). The RSMD of the ExoN polypeptide chain progressively stabilizes during the course of the simulation, and better stabilization of the protein was observed in the presence of RNA (**Sup Fig 7a**). RMS fluctuations at the Cα positions across the polypeptide chain indicate that the protein demonstrates extensive movement at the N-terminal NSP10 binding region, whereas ZnF as well as core-catalytic region showed minimal fluctuations of the order of 0.1 nm(**Sup Fig 7b**). Based on binding energy estimates, ZnF1 appears to bind single stranded RNA and ZnF2 binds double stranded RNA with substantial affinity (**Sup Fig 7c**). Major residues involved in binding were identified based on the residence time of their interaction with RNA, which could be mapped to ZnF1 and 2 (**Fig 3b & 3d**, **Sup Fig 6b & 6d**). Thereby, this viral exonuclease appears to exploit the RNA binding ability of the ancestral ZnFs for RNA recognition for proofreading.

**Figure 3:**
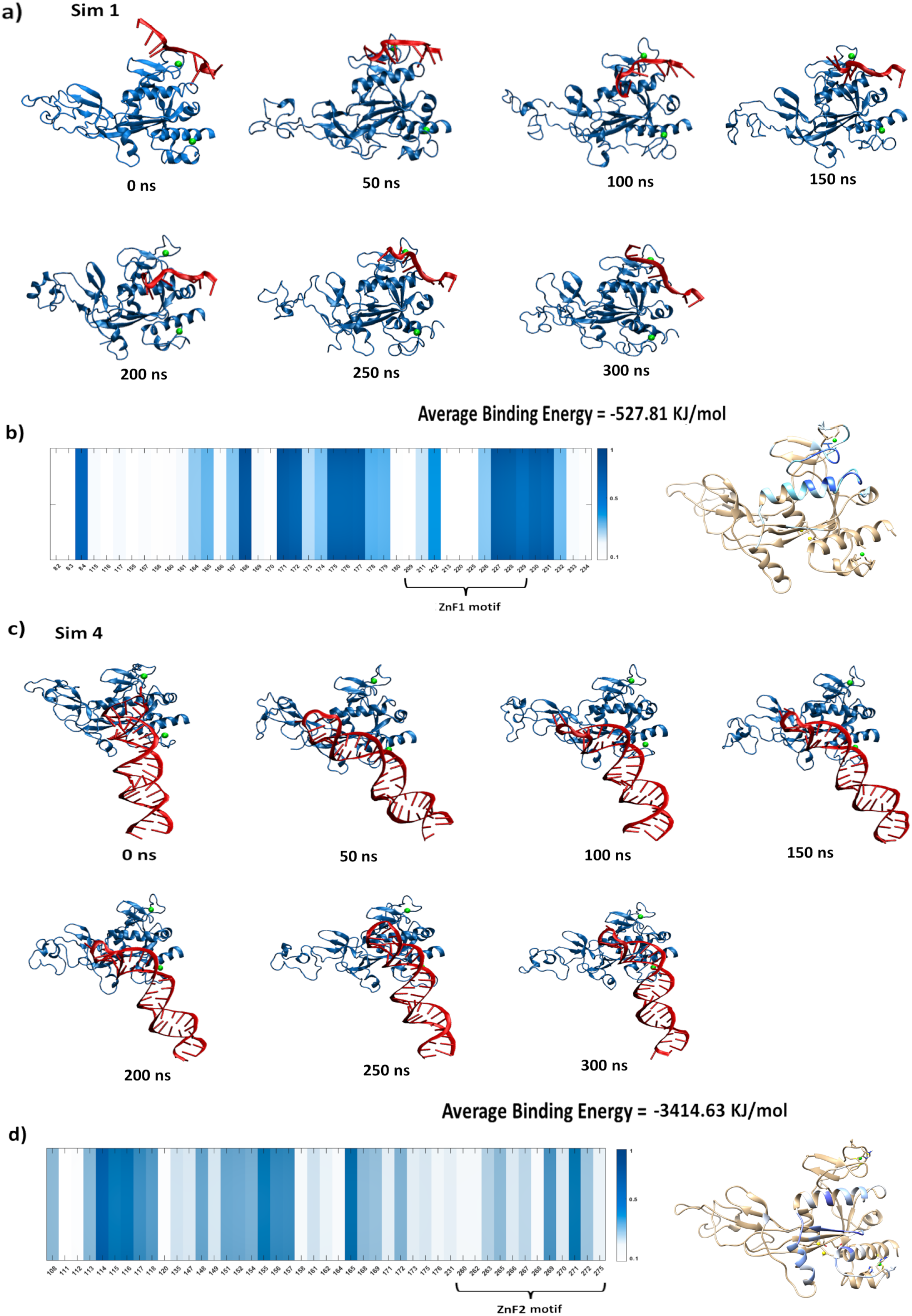
Molecular Dynamics simulations identifies RNA binding to ZnFs in ExoN. **a) & c) Dynamics of RNA interaction with ZnF1 and ZnF2 of ExoN** The dynamics of single stranded RNA interaction with ZnF1 (Fig 3a) and double stranded RNA with ZnF2 (Fig 3c) at every 50ns time points is represented. The ExoN is depicted in blue cartoon. RNA is depicted in red cartoon. Zinc ions are represented in green. The average binding energy between RNA and ExoN estimated from the first 150ns simulation is represented at the bottom. **b) & d) ExoN residues involved in RNA interaction** The residues of ExoN interacting with RNA throughout the simulation is represented as a heatmap. The color scale depicts the fraction of time during which the residue contacts RNA moiety. The residues involved in the RNA interaction are mapped on to the ExoN structure and color coded based on residence time with the same scale bar.

### Structural comparison of ExoN with DNA proofreading machinery suggests convergent evolution

Proofreading DNA polymerases and ExoN share structural similarity. DALI search identified exonuclease domains of *E. coli* DNA polymerase 1 and 2, as well as the epsilon subunit of DNA polymerase 3 as significant hits that are also biochemically well studied. While the catalytic core was similar in all these proofreaders, the entire ZnF1 motif was unique to ExoN. Though the ZnF2 motif was not present in DNA proofreaders, helices that harbor zinc chelating residues showed substantial superposition (**Fig 4a**). ZnF1 motif with a beta hairpin and loops harboring zinc chelating residues (CCCH) were distinct, and resembled the beta-hairpin in DNA polymerase 2 involved in mismatch recognition(Wang and Yang 2009) suggesting a similar role for ZnF1 (**Fig 4b**). Substantiating this possibility, the intermediate sized genomes of mesoniviridae and medioniviridae lacked a key ZnF1 residue involved in zinc chelation (**Sup Fig 3b & Fig 4c**). Suggesting that these intermediate size genomes could have arisen due to compromised proofreading ability contributed by the ZnF1.

**Figure 4:**
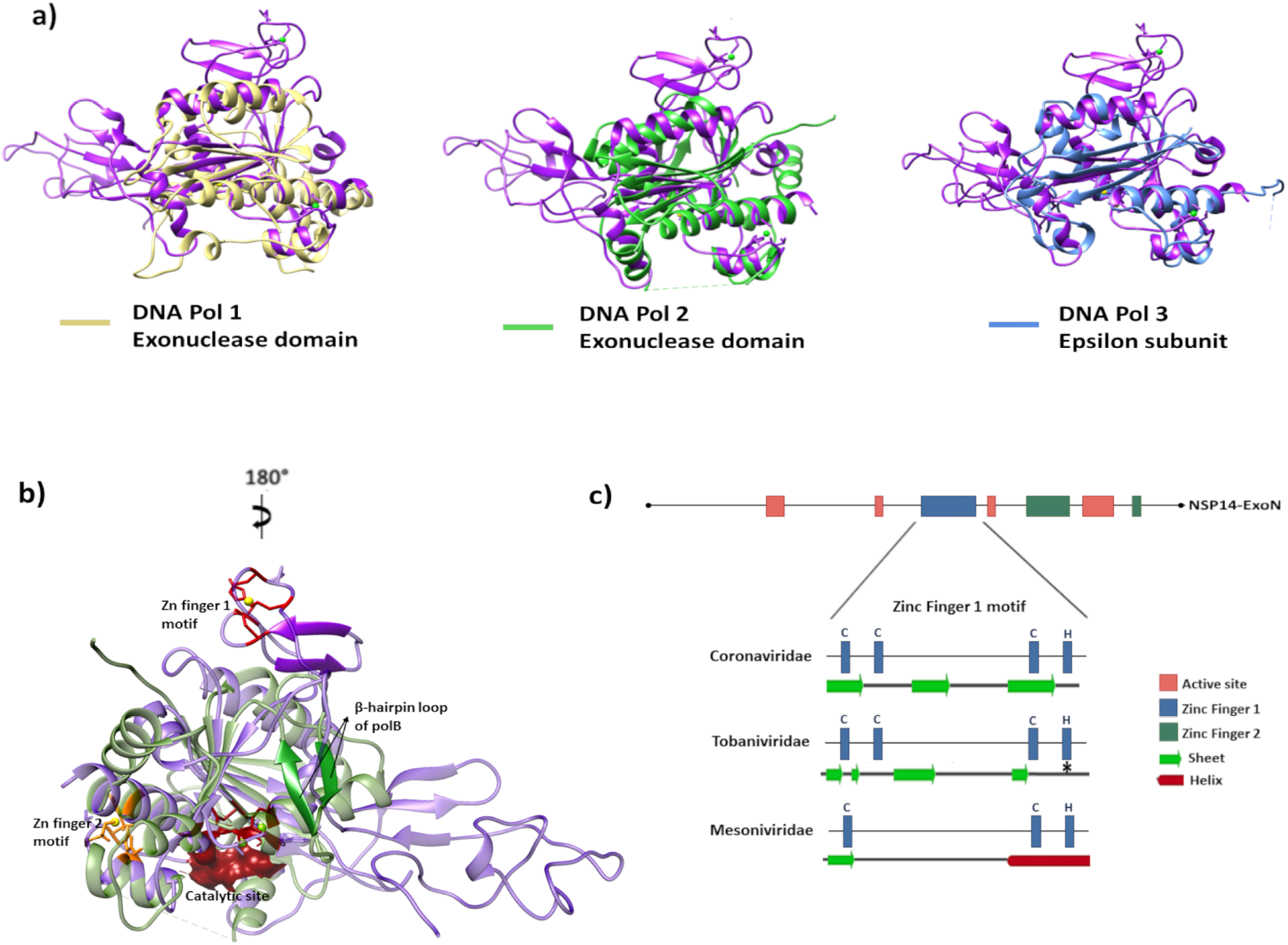
Structural comparison of ExoN with DNA proofreaders. **a) Structural superpositioning** Structural superpositioning of ExoN (in purple) with Exonuclease domain of *E. coli* DNA Pol I (in yellow), *E.coli* DNA Pol II (in green) and epsilon subunit of *E. coli* Pol III (in blue) is depicted. **b) Superimposition of DNA polymerase 2 exonuclease domain with that of SARS-CoV2 NSP14.** Purple color represents SARS-CoV2 NSP14 exonuclease domain while green color represents DNA polymerase 2. Zinc ions are represented in yellow and Magnesium ions are represented in light green. Zinc finger 1 residues are represented in red color and Zinc finger 2 residues are represented in orange. Catalytic site residues are represented as surface in brown color. Beta-hairpin motif of DNA Pol II involved in DNA recognition is highlighted in green and the beta-hairpin in the ZnF1 in purple. **c) Secondary structure prediction in the Zinc finger 1 motif** Sequence-based secondary structure prediction around the ZnF 1 motif for Coronaviridae, Mesoniviridae and Tobaniviridae are represented. The * sign represents that the residue is not conserved among all the sequences analyzed.

Proofreading function is an elaborate multiprotein activity involving mismatch sensing by the polymerase, unwinding of the duplex by helicase, confirmation of the mismatch by exonuclease followed by 3’ to 5’ cleavage. Given their ability to bind RNA, ZnFs may have been co-opted for mismatch recognition and led to this convergent mode of proofreading. The pattern of codon usage indicated a gradual adaptation of NSP14 in nidovirales, having achieved a synchronized pattern in coronaviral genome (**Fig 2d**). Analyzing the pattern of amino acid substitutions within the nidovirales suggests that polyprotein cleaving protease (NSP3) to highly coevolve with almost all NSPs encoded (**Sup Fig 8**). While ExoN formed a cluster with replicase and helicase, it was only moderately coevolving. However, on closer analysis we observed that a key residue involved in catalysis and the two zinc finger motifs demonstrate synchronous substitutions suggesting coevolution with several NSPs. Maximal coevolution was observed with NSP12 and NSP13 involved in replication and helicase function respectively (**Fig 5 a & b**). Thereby we propose that these viruses have acquired a rudimentary exonuclease with a capacity to recognize RNA using the Zinc finger motifs that has convergently evolved to function as a proofreading enzyme by coordinating with the core replication complex.

**Figure 5:**
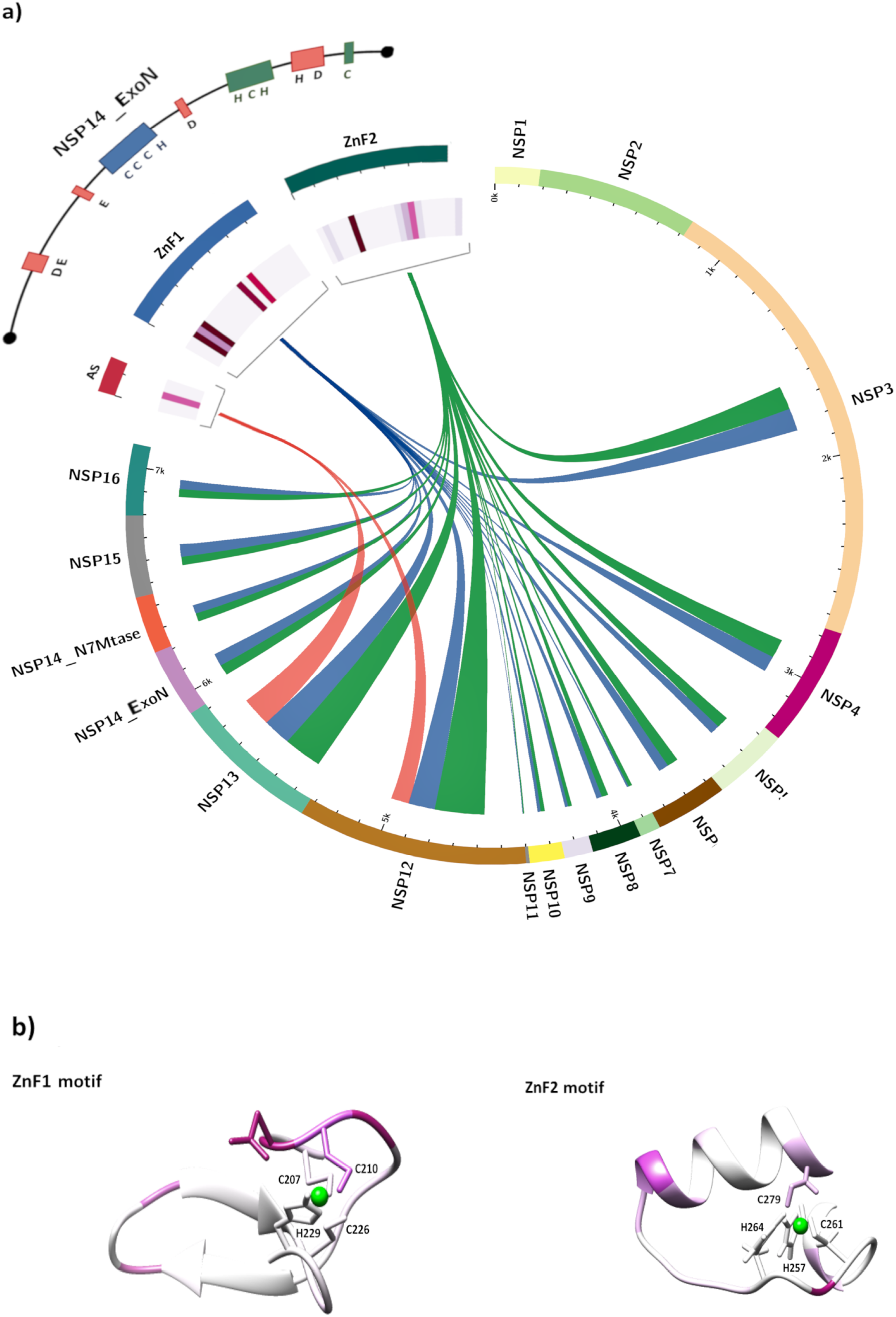
Coevolution of ExoN active site residues and ZnF motifs. **a) Synchronous substitution in ExoN with NSP12 and 13 suggest coevolution of proof-reading** Coevolution of ExoN active site residues and both ZnF motifs with all the other NSPs in the genome are visualized as a circos plot. Connectivity depicts residues in active site (DEDDh), ZnF1 or ZnF2 motifs that coevolve with each NSP and is represented in red, blue and green ribbons respectively with width proportional to percentage of coevolving residues. The heatmap represents residue wise cumulative coevolution across NSP12 and 13 depicted for active site, ZnF1 and ZnF2 towards the top left. **b) Structural mapping of coevolution in ZnF1 and ZnF2 motif** The heatmap represented in Fig 4a depicting the total number of residues with NSP12 and 13, that coevolve with each residue in ZnF1 and 2 motifs are mapped to structure of ExoN.

## Discussion

RNA proofreading is a challenge due to the relative stability of local structures in RNA than its DNA counterpart. Hence these viruses seemingly have evolved a complex mechanism to recognize the RNA mismatch for proofreading that differs from cellular organisms as well as DNA viruses. Deployment of DEDDh family exonuclease though is common, use of zinc fingers to recognize the replicating genome is a novel feature of RNA proofreaders, which could be a complex but necessary means to recognize RNA mismatch. A large RNA genome due to its proof-reading ability has seemingly permitted the acquisition of newer enzymes. Recently, the role ExoN in recombination of SARS-CoV2 has emerged, which could have independently facilitated lateral gene transfer events(Gribble et al. 2021). Consequently, the complex genomes of coronavirus are likely to enable self-sustenance of these viruses and their reliance on the host machinery is reduced. It remains to be seen whether this could permit zoonotic spillover of coronaviruses by fewer adaptive mutational events. Nature has seemingly experimented with the acquired enzyme to optimize proofreading and balance evolvability with genome size. This is evident in the tinkering of zinc fingers which are critical to couple RNA binding to exonuclease activity. While the medium sized genomes of mesoniviridae and medioniviridae are likely to have intermediate level of proofreading capability due to destabilized ZnFs, larger genomes of coronaviridae have further fine-tuned the zinc fingers to coevolve with the core machinery and arrive at optimal level of proofreading. Thereby in this study by tracing the evolutionary history of this protein family, we provide molecular correlates for the breach of Eigen’s paradox.

Multiple extinction events of H1N1 influenza virus in the past century have been documented(Carter and Sanford 2012). This is largely attributable to its high mutation rate and inability to sustain a stable genome. In contrast, the relative low mutation burden of SARS-CoV2 would make such genetic attenuation a weak possibility and lead to waves of localized infection due to escape variants that are currently being experienced by several countries. Hence, vaccinations and therapeutics in combination, hold the only hope for managing the current pandemic. Ironically, the most promising antiviral Remdesivir and allied nucleoside analogs are required in high dosages because of exonucleolytic removal from RNA by ExoN, that decreases their efficacy(Agostini et al. 2018; Ferron et al. 2018; Robson et al. 2020). Hence, Remdesivir in combination with ZnF-targeting ExoN inhibitor would be a good approach to improve efficacy. However, this would pose additional challenge of increasing the mutation rate and could even result in the emergence of more virulent strains. Alternately, we propose that an allosteric activator of ExoN, binding to ZnF1/2 would be a better strategy to self-destruct the growing RNA and curtail viral replication.

## Online Methods

### Collection of sequences

All RNA viruses having the host specificity to human and viruses belonging to Nidovirales order were identified from ViralZone Expasy(Viralzone) website and further validated using International Committee on Taxonomy of Viruses (ICTV)(Lefkowitz et al. 2018) website. The protein sequences of the non-structural proteins of all the identified viruses were obtained from the Protein database of the NCBI RefSeq(Pruitt et al. 2005). Their total genome size, individual ORF size were also obtained from NCBI.

### Pairwise alignment

For the genomes that are not well characterized and annotated, the respective NSPs were identified using pairwise sequence alignment using EMBOSS-NEEDLE(Madeira et al. 2019) webserver. Also, to identify the similarity between two sequences, pairwise alignment was performed using Needleman-Wunsch alignment algorithm with the help of EMBOSS-NEEDLE webserver.

### SARS-CoV2 Variant’s data analysis

The variation data from all the sequenced SARS-CoV2 genomes were extracted from China National Center for Bioinformation (https://bigd.big.ac.cn/ncov)(Gong et al. 2020; Song et al. 2020; Zhao et al. 2020) till February 5^th^2021.

### H1N1 Influenza Variant’s data analysis

Data was obtained from the NIAID Influenza Research Database (IRD)(Zhang et al. 2017) through the web site (http://www.fludb.org). For comparable dataset, H1N1 human Influenza virus variations was collected from January 2009 to June 2010 pandemic by running SNP analysis in the fludb server.

### Visualization of variation data

The mutation burden in each gene was calculated as the number of base pairs in the gene (having mutation frequency more than 0.01) divided by the gene size. SNP mutation frequency across the genome was plotted as circus plot using the Circos package v0.69-9(Krzywinski et al. 2009).

### Multiple Sequence Alignment

The webserver MAFFT (Multiple Alignment using Fast Fourier Transform)(Katoh et al. 2002) was used to construct multiple sequence alignment of the sequences under study.

### Phylogenetic tree/ cladogram construction

The aligned NSP-12 sequences was preprocessed to the required PHYLIP format using JALVIEW v2.11.1.3(Waterhouse et al. 2009). The phylogenetic tree construction was performed using raxmlHPC module of raxml-7.3.0-release(A.P. Stamatakis et al. 2008) under GTR AA substitution model with estimate of proportion of invariable sites and auto rapid bootstrap convergence algorithm(A. Stamatakis et al. 2008). The tree was visualized using Figtree v1.4.4(Rambaut 2009). Zinc finger proteins that bind to RNA were identified from literatures. The sequences of well characterized Zinc finger motifs from these proteins are collected and used along with Zinc fingers in SARS-CoV2 ExoN to construct a phylogenetic tree similar to the methodology explained above.

### Pinpointing the ExoN origin by Most Common Recent Ancestor analysis

Position Specific Iterative-Blast (PSI-BLAST)(Altschul and Koonin 1998) was performed using PSI-BLAST standalone tool(Madden 2013) to find out the distant orthologs. The sequences of Tobaniviridae family were used as the query and distant orthologs were searched in Refseq protein database with default settings. The iterations were performed till the algorithm could not find any new sequences. Convergence of iterations was achieved at iteration 23, and the hits were sorted based E-Value and total score. Also, the top potential hits were analyzed for the presence of DEDDh residues and Zinc finger residues.

### Codon usage

The codon usage pattern of NSP12 and NSP14 ExoN in Tobaniviridae, Mesoniviridae and Coronaviridae was calculated using EMBOSS Cusp webserver(Rice et al. 2000) and visualized as a heatmap in RStudio. Chinook salmon virus, Nam Dinh virus and SARS-CoV2 were taken as representatives of Tobaniviridae, Mesoniviridae and Coronaviridae respectively. The Euclidean distance between NSP12 and NSP14 ExoN was calculated using dist function in RStudio.

### Structure modelling and validation

The structure of SARS-CoV2 NSP14 was modelled based on SARS-CoV NSP14 (PDB #5C8S) using I-TASSER web server(Zhang 2008). The model validation was performed by structure evaluation servers like ERRAT(Petcov 1977), WHATCHECK(Vriend 1990), PROCHECK(Laskowski et al. 1993), Prove(Pontius et al. 1996) and Verify3D(Eisenberg et al. 1997).

### Binding Pocket Prediction

The Modelled SARS-CoV2 NSP14 structure was used to predict binding pockets in ExoN structure via CASTp webserver(Tian et al. 2018). The top 3 binding pockets in exonuclease domain having higher SASA volume was analyzed. Out of the three pockets, one of them was present near the ZnF1 motif, second in the Active site motif. These three pockets were also predicted by other major tools such as fpocket(Le Guilloux et al. 2009) and prank(Krivák and Hoksza 2015; Krivák and Hoksza 2018).

### Docking

Two binding pockets present near the ZnF1 motif and the Active site motif was used to define the active residues for docking of both ssRNA and dsRNA in Haddock docking tool(De Vries et al. 2010). A docking of NSP14 with ssRNA and dsRNA was also performed using Prime-3D2D docking tool(Xie et al. 2020) and the RNA docked majorly to ZnF2 and occupied little near ZnF1. The docked structures from these four different modes were used for further simulation studies. The structure of ssRNA (with a sequence 5’-UUAUUUAUU-3’) was taken from PDB #1RGO and dsRNA (with a sequence, Primer: 5’-CAUGCUACGCGGUAGUAGCAUGCUAGGGAGCAG-3’, Template: 5’-CAUGCCAUGGCCUGUAAAAUGUCUGACUGCUCCCUAGCAUGCUACUACCGCGUAGCAUG-3’) structure was taken from PDB #7CYQ

### Molecular Dynamics Simulation

Each of the four docked structures were used as starting conformations, placed in a cubic box and solvated using TIP3P(Mark and Nilsson 2001) water model. All the systems were equilibrated using the standard minimization and equilibration steps. Simulations were carried out on all the systems using GROMACS(Van Der Spoel et al. 2005) by employing AMBER all-atom force field. Periodic boundary conditions were used and 1.2nm was set as real space cut-off distance. Particle Mesh Ewald (PME) summation with grid spacing of 0.16 nm was used combining with a fourth-order cubic interpolation to deduce the forces and potential in-between grid points. The van der Waals cut off was set to 1.0nm. A 2fs time step for numerical integration of the equations of motion was used and the coordinates were saved at every 100ps. The initial systems were subjected to energy minimization using the steepest descent method. Modified Berendsen thermostat T-coupling bath was used to maintain 310K temperature and Parrinello-Rahman barostat was used for pressure coupling at 1 bar, before starting the 300 ns production run. The Zinc coordination between cysteine and histidine residues in ZnF1 and ZnF2 motif was defined in the simulation based on the parameters file from Macchiagodena et al(Macchiagodena et al. 2019).

### Analysis of Trajectories

The global analysis of trajectories was carried out by computing Root Mean Square Deviation (RMSD) and Root Mean Square Fluctuation (RMSF) using GROMACS modules rms and rmsf respectively. The interaction between the protein and RNA throughout the simulation time was calculated using MDCons package(Abdel-Azeim et al. 2014). The binding energy of the RNA with protein was calculated using g_mmpbsa package(Kumari et al. 2014). Graphing, Advanced Computation and Exploration (GRACE) program, version 5.1.22 and MATLAB were used to construct graphs/plots. VMD(Humphrey et al. 1996) and UCSF Chimera(Pettersen et al. 2004) were used for visualization and generation of molecular images.

### Structure analysis

DALI(Holm and Sander 1995) search was performed to identify the crystal structures of proof-reading DNA polymerases having structural similarity to SARS-CoV NSP14 structure (PDB #5C8S). The hits from *E. coli* DNA polymerases included DNA polymerase 1 (z, 8.2), DNA polymerase 2 (z, 8.2) and DNA polymerase 3 epsilon subunit (z, 10.2). Structural superposition of these hits with NSP14 was performed in chimera. The secondary structure prediction in the ZnF1 motif for Tobaniviridae, Mesoniviridae and Coronaviridae was done using Jpred tool(Cuff et al. 1998) in JALVIEW software(Waterhouse et al. 2009). The predicted structure of human ZNFX1 was obtained from AlphaFold protein structure database(Jumper et al. 2021).

### Coevolution analysis

The coevolution between the residues of SARS-CoV2 polyprotein 1ab was calculated as a chi-square statistic covariance metric using CoeViz web-based tool(Baker and Porollo 2016) integrated in Polyview-2D server(Porollo et al. 2004). The residues that had covariance score less than 0.6 were filtered out. The percentage of residues in each protein coevolving with the other was calculated and plotted as a heatmap. The number of residues in each protein coevolving with the active site, ZnF1 and ZnF2 motifs of NSP14_ExoN were calculated and plotted as a circos plot using Circos package v0.69-9. The number of residues in NSP12 and NSP13 coevolving with individual residues in the active site, ZnF1 and ZnF2 motifs of NSP14_ExoN were calculated and represented as a heatmap in the circos. This heatmap was also depicted in the structure of ZnF1 and ZnF2 motifs using Chimera software to map coevolving residues between ZnFs and core replication machinery.

The heatmaps were generated using Complex Heatmap package(Gu et al. 2016) in Rstudio v1.3.959(Allaire 2012) and boxplots were also generated in RStudio. High quality images were generated using GIMP (GNU image manipulation program) v2.10(team 2013)

## Acknowledgements

This work was supported by the Department of Biotechnology, India through grant (GAP0214, BT/PR40396/COT/142/7/2020) and Council for Scientific and Industrial Research (CSIR), India through grant (MLP2008) for the support of infrastructure and fellowship to SKA. CSIR-4PI for supercomputing facilities. Dr. Lipi Thukral (CSIR-Institute of Genomics and Integrative Biology) and Dr. Vineet Gaur (National Institute of Plant Genome Research) for help with molecular dynamics simulations and structure interpretations respectively.

## Author Contributions

SKA and VTN conceptualized, SKA executed the analyses and was supervised by SR and VTN. SKA and VTN interpreted the data and wrote the manuscript.

## Supplementary Figure legends

**Supplementary Figure 1:**

**Percentage of Virus families with predicted Zoonotic spillover potential**

The percentage of viruses from different families with predicted zoonotic spillover potential are represented as a pie chart. Key families and their causative viruses are highlighted along with their disease manifestation.

**Supplementary Figure 2:**

**a) Functionally characterized Non-structural Protein comparison among Nidovirales order.**

Matrix representing the presence or absence as black and while squares for non-structural proteins of different families in the Nidovirales order. The median of the genome size for each family is plotted as violet bar plot to the right.

**b) Viral genome size across groups of viruses that either contain proof-reading machinery or lack ExoN (+ /−) among the Nidovirales order**

Box plots represent the size in kb of the whole genome, ORF1A, ORF1B and other 3’ ORFs of viruses in all families relative to the presence and absence of ExoN proof-reading machinery. Horizontal line represents median value and whiskers represent 4^th^ and 1^st^ quartile at top and bottom respectively.

**c) Genome size comparison among Nidovirales order**

Box plot depicts the size in kb of the whole genomes representing indicated family of viruses in nidovirales order. Horizontal line represents median value and whiskers represent 4^th^ and 1^st^ quartile at top and bottom respectively and categorized into small ≤ 16000 bp, large ≥ 25000 bp and intermediate sizes of around 20000 bp.

**Supplementary Figure 3:**

**a) Multiple Sequence Alignment of ExoN orthologs and DEDDh family exonucleases**

Representative members of Nidovirales ExoN along with proofreading exonucleases from bacteria and exonucleases from plant, invertebrate and vertebrate species were subjected to multiple sequence alignment and the region pertaining to the active site residues DEDDh is depicted. The red boxes represent the catalytic sites, blue boxes represent zinc finger 1 (ZnF1) residues and green boxes represent zinc finger 2 (ZnF2) residues.

**b) Architecture of ExoN**

Domain architecture from all families of Nidovirales representing residues involved in catalysis (DEDDh), ZnF 1 and 2 is depicted.

**c) Identification of Common Recent Ancestor of ExoN**

Output after convergence of Position Specific Iterative (PSI) Blast of Tobaniviridae ExoN is depicted as a table.

**Supplementary Figure 4:**

**Tracing the ligand specificity of ZnF based on phylogeny**

Well-characterized Zinc finger domains and their orthologs were subjected to multiple sequence alignment along with ZnF1 and 2 of ExoN and the cladogram is represented along with the RNA specificity.

**Supplementary Figure 5:**

**a) Modelling and validation of SARS-CoV2 NSP14**

The structure of SARS-CoV2 was modelled based on SARS-CoV NSP14 crystal structure and the superposition of the backbone is depicted. RMSD parameters wrt SARS-CoV and the recently available ExoN of SARS-CoV2 structures is depicted below. Table represents quality control parameters for validating the structure for subsequent studies.

**b) Prediction of binding pocket**

Output of binding pocket prediction in ExoN is depicted.

**c) Docking analysis of RNA**

Single and double stranded RNA is docked at both ZnF1 and 2 and the combinations are used for Molecular Dynamic simulations to assess binding.

**Supplementary Figure 6:**

**a) & c) Dynamics of RNA interaction with ZnF1 and ZnF2 of ExoN**

The dynamics of double stranded RNA interaction with ZnF1 (Sup Fig S7a) and single stranded RNA with ZnF2 (Sup Fig S7c) at every 50ns time point is represented. The ExoN is depicted in blue cartoon and RNA in red cartoon. Zinc ions are represented in green. The average binding energy between RNA and ExoN estimated from the first 150ns simulation is represented at the bottom.

**b) & d) ExoN residues involved in RNA interaction**

The residues of ExoN interacting with RNA throughout the simulation is represented as a heatmap. The color scale depicts the fraction of time during which the residue contacts RNA moiety. The residues involved in the RNA interaction are mapped on to the ExoN structure and color coded based on residence time with the same scale bar.

**Supplementary Figure 7:**

**a) RMS deviation plot for the ExoN during the course of simulations**

**b) RMS fluctuations at the Cα positions for the residues in ExoN during the course of simulations.**

**c) Binding energy of the RNA-ExoN interactions calculated as ΔG during the first 150ns of simulations.**

**Supplementary Figure 8:**

Coevolution of residues within the Nidovirales genome was calculated based on synchronous substitution in pairwise comparison of aminoacids across the NSPs. Percent residues within a protein coevolving with the rest of the NSPs is depicted as a heatmap, which is clustered hierarchically. The scale bar for the heat map is depicted.

**Supplementary Table 1**

Accession number of the viral genome sequences used for the analysis and the accession number for Zinc Finger Proteins are provided in a tabular form.

