## Supplementary Figures for "Adaptive convergent evolution of genome proofreading in SARS-CoV2: insights into the Eigen’s paradox"

Sup Fig 1

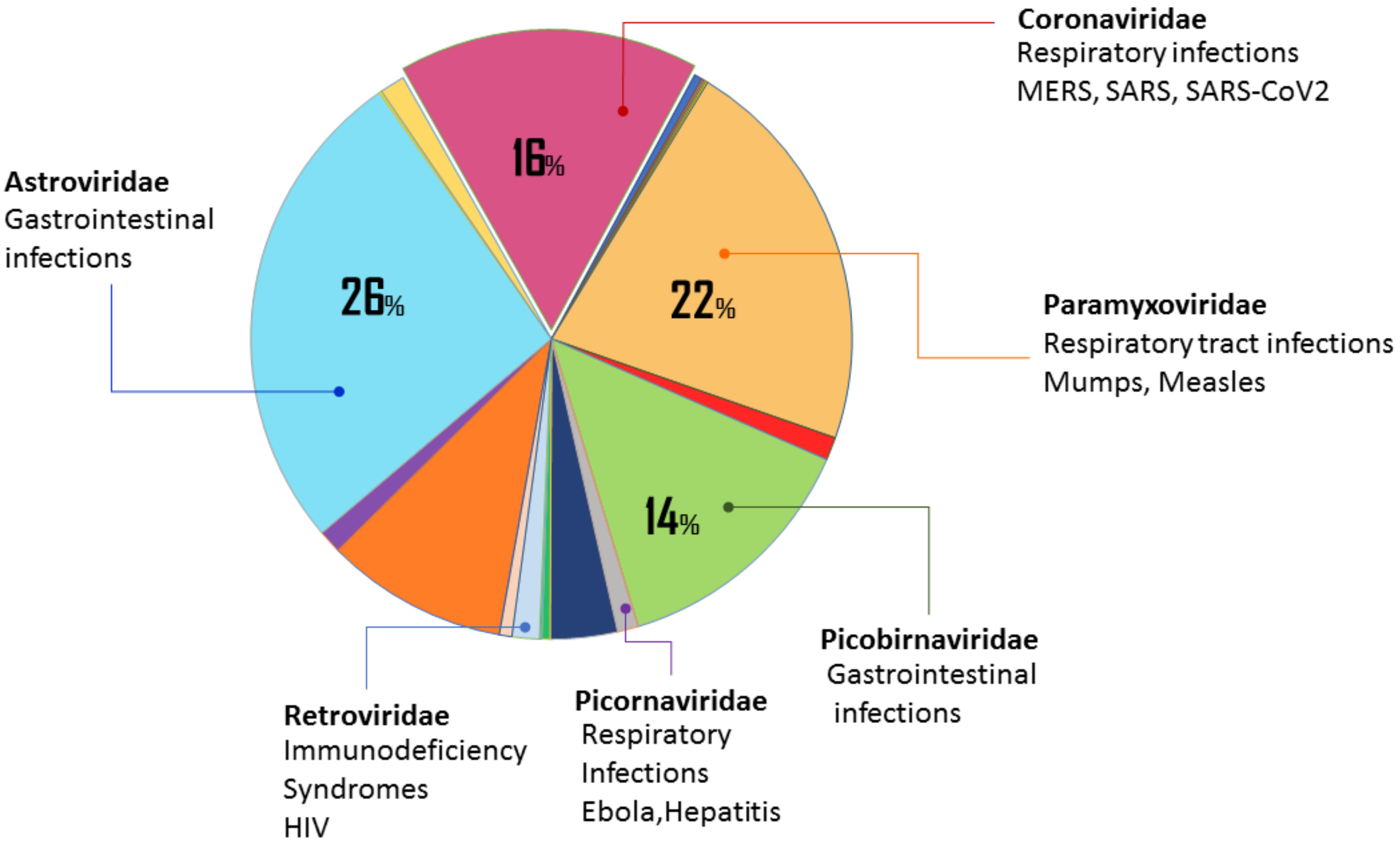
16%
22%
26%
14%
Coronaviridae
Respiratory infections
MERS, SARS, SARS-CoV2
Astroviridae
Gastrointestinal
infections
Paramyxoviridae
Respiratory tract infections
Mumps, Measles
Picobirnaviridae
Gastrointestinal
infections
Retroviridae
Immunodeficiency
Syndromes
HIV
Picornaviridae
Respiratory
Infections
Ebola,Hepatitis

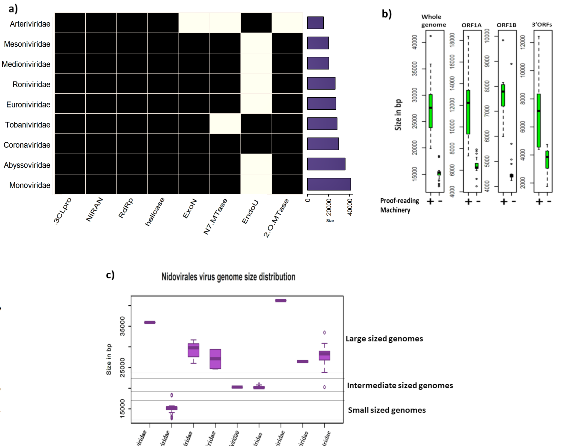

a)
Arteriviridae
Mesoniviridae
Medioniviridae
Roniviridae
Euroniviridae
Tobaniviridae
Coronaviridae
Abyssoviridae
Monoviridae
3CLpro
NiRAN
RdRp
helicase
ExoN
N7.MTase
EndoU
2.O.MTase
0
20000
40000
Size
b)
Whole genome
ORF1A
ORF1B
3'ORFs
Size in bp
Proof-reading Machinery
+
-
c)
Nidovirales virus genome size distribution
Size in bp
35000
25000
15000
Large sized genomes
Intermediate sized genomes
Small sized genomes
Abyssoviridae
Arteriviridae
Coronaviridae
Euroniviridae
Medioniviridae
Mesoniviridae
Monoviridae
Roniviridae
Tobaniviridae


Sup Fig 2

### Sup Fig 3

a)

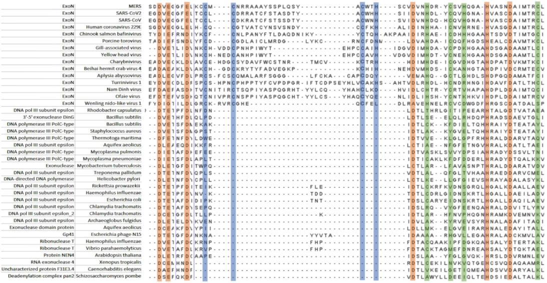

b)

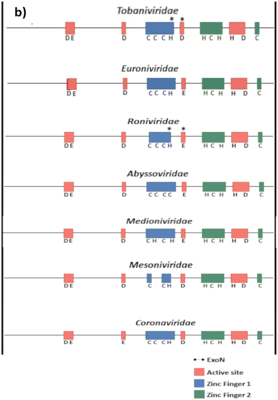


c)

| Protein | Species | Phylum | Total Score | Query Cover | E value | Per. ident | Accession |
|---|---|---|---|---|---|---|---|
| NFX1-type zinc finger-containing protein 1-like | Strongylocentrotus purpuratus | Echinodermata | 1351 | 99% | 0 | 10.4 | XP_030846999.1 |
| PREDICTED: cyclin-D1-binding protein 1 | Cuculus canorus | Chordata | 823 | 99% | 0 | 10.77 | XP_009556686.1 |
| hypothetical protein LOTGIDRAFT_213601 | Lottia gigantea | Mollusca | 748 | 99% | 2.00E-176 | 12.39 | XP_009050447.1 |
| hypothetical protein HELRODRAFT_65033 | Helobdella robusta | Annelida | 502 | 97% | 1.00E-159 | 11.43 | XP_009015652.1 |
| hypothetical protein GGTG_13333 | Gaeumannomyces tritici | Ascomycota | 612 | 98% | 1.00E-146 | 10.92 | XP_009229503.1 |
| uncharacterized protein FOYG_05943 | Fusarium oxysporum | Ascomycota | 765 | 98% | 7.00E-146 | 10.11 | XP_031041286.1 |
| hypothetical protein ASPGLDRAFT_46703 | Aspergillus glaucus | Ascomycota | 783 | 98% | 4.00E-145 | 8.76 | XP_022401491.1 |
| hypothetical protein TERG_02292 | Trichophyton rubrum | Ascomycota | 661 | 99% | 4.00E-144 | 11.05 | XP_003237575.1 |
| hypothetical protein COCC4DRAFT_62859 | Bipolaris maydis | Ascomycota | 754 | 99% | 6.00E-143 | 10.74 | XP_014076953.1 |
| hypothetical protein SAMD00019534_118060 | Acytostelium subglobosum | Amoebozoa | 595 | 98% | 4.00E-142 | 8.99 | XP_012748408.1 |
| uncharacterized protein SETTUDRAFT_96932 | Exserohilum turcica | Ascomycota | 633 | 98% | 6.00E-141 | 10.88 | XP_008030358.1 |
| uncharacterized protein THITE_51448 | Thermothielavioides terrestris | Ascomycota | 838 | 99% | 4.00E-139 | 9.02 | XP_003654692.1 |
| hypothetical protein BO94DRAFT_563445 | Aspergillus sclerotioniger | Ascomycota | 747 | 98% | 1.00E-138 | 9.98 | XP_025470651.1 |
| uncharacterized protein BDV37DRAFT_292935 | Aspergillus pseudonomius | Ascomycota | 628 | 99% | 3.00E-138 | 8.16 | XP_031943062.1 |
| hypothetical protein NEUTE1DRAFT_68071 | Neurospora tetrasperma | Ascomycota | 733 | 99% | 9.00E-138 | 9.28 | XP_009853737.1 |
| hypothetical protein BO85DRAFT_523897 | Aspergillus piperis | Ascomycota | 745 | 98% | 1.00E-137 | 9.23 | XP_025510987.1 |
| hypothetical protein PV07_12556 | Cladophialophora immunda | Ascomycota | 545 | 99% | 1.00E-137 | 10.24 | XP_016242267.1 |
| P-loop containing nucleoside triphosphate hydrolase protein | Phialocephala scopiformis | Ascomycota | 716 | 99% | 3.00E-137 | 8.61 | XP_018064031.1 |
| hypothetical protein PgNI_06085 | Pyricularia grisea | Ascomycota | 880 | 99% | 7.00E-137 | 9.36 | XP_030982267.1 |
| hypothetical protein DICPUDRAFT_147315 | Dictyostelium purpureum | Amoebozoa | 439 | 96% | 1.00E-135 | 8.83 | XP_003283594.1 |
| Pc13g10110 | Penicillium rubens | Ascomycota | 821 | 99% | 1.00E-134 | 10.88 | XP_002559433.1 |
| similar to NF-X1 finger and helicase domain protein | Leptosphaeria maculans | Ascomycota | 847 | 100% | 2.00E-134 | 10.8 | XP_003835891.1 |
| DNA-binding protein smubp-2, putative | Talaromyces stipitatus | Ascomycota | 664 | 98% | 3.00E-123 | 10.86 | XP_002484253.1 |
| uncharacterized protein LOC117289197 | Asterias rubens | Echinodermata | 1754 | 99% | 2.00E-122 | 11.78 | XP_033626096.1 |
| conserved hypothetical protein | Phytophthora infestans | Oomycota | 574 | 0.97 | 1.00E-121 | 9.15 | XP_002902248.1 |

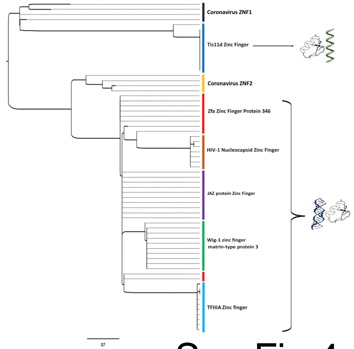

Coronavirus ZNF1
Tis11d Zinc Finger
Coronavirus ZNF2
Zfa Zinc Finger Protein 346
HIV-1 Nucleocapsid Zinc Finger
JAZ protein Zinc Finger
Wig-1 zinc finger
matrin-type protein 3
TFIIIA Zinc finger
0.7


Sup Fig 4

**a)**

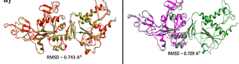


I-Tasser model - Green color
SARS Coronavirus nsp-14 - Red color

I-Tasser model - Green color
SARS-CoV2 NSP14_ExoN – Magenta color

| MODEL | C-Score | Verify 3D Score | ERRAT Quality Factor | Ramachandran Plot (residues in disallowed regions) |
|---|---|---|---|---|
| I-Tasser model | 2 | 87.67% | 90.02 | 1.1% |

**b)**

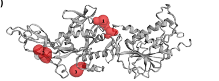


### Sup Fig 5

| Pocket Number | Area (SA) | Volume (SA) |
|---|---|---|
| 1 | 228.497 | 108.027 |
| 2 | 94.882 | 73.967 |
| 3 | 87.130 | 68.175 |

**c)**

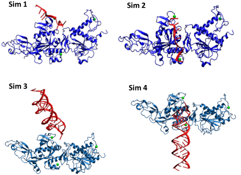

Sup Fig 6

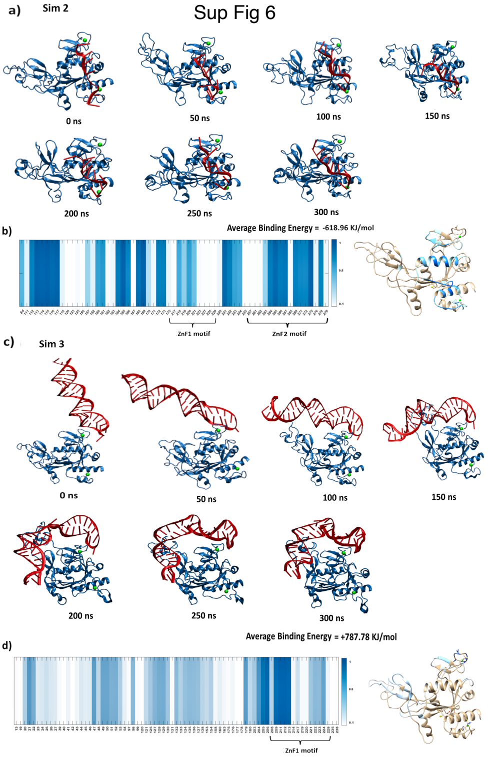

a)
Sim 2
0 ns
50 ns
100 ns
150 ns
200 ns
250 ns
300 ns
b)
Average Binding Energy = -618.96 KJ/mol
ZnF1 motif
ZnF2 motif
c)
Sim 3
0 ns
50 ns
100 ns
150 ns
200 ns
250 ns
300 ns
d)
Average Binding Energy = +787.78 KJ/mol
ZnF1 motif

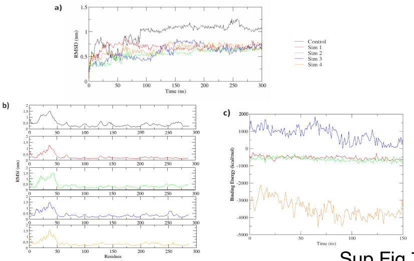
a)
RMSD (nm)
Time (ns)
Control
Sim 1
Sim 2
Sim 3
Sim 4
b)
RMSF (nm)
Residues
c)
Binding Energy (kcal/mol)
Time (ns)

Sup Fig 7

### Sup Fig 8

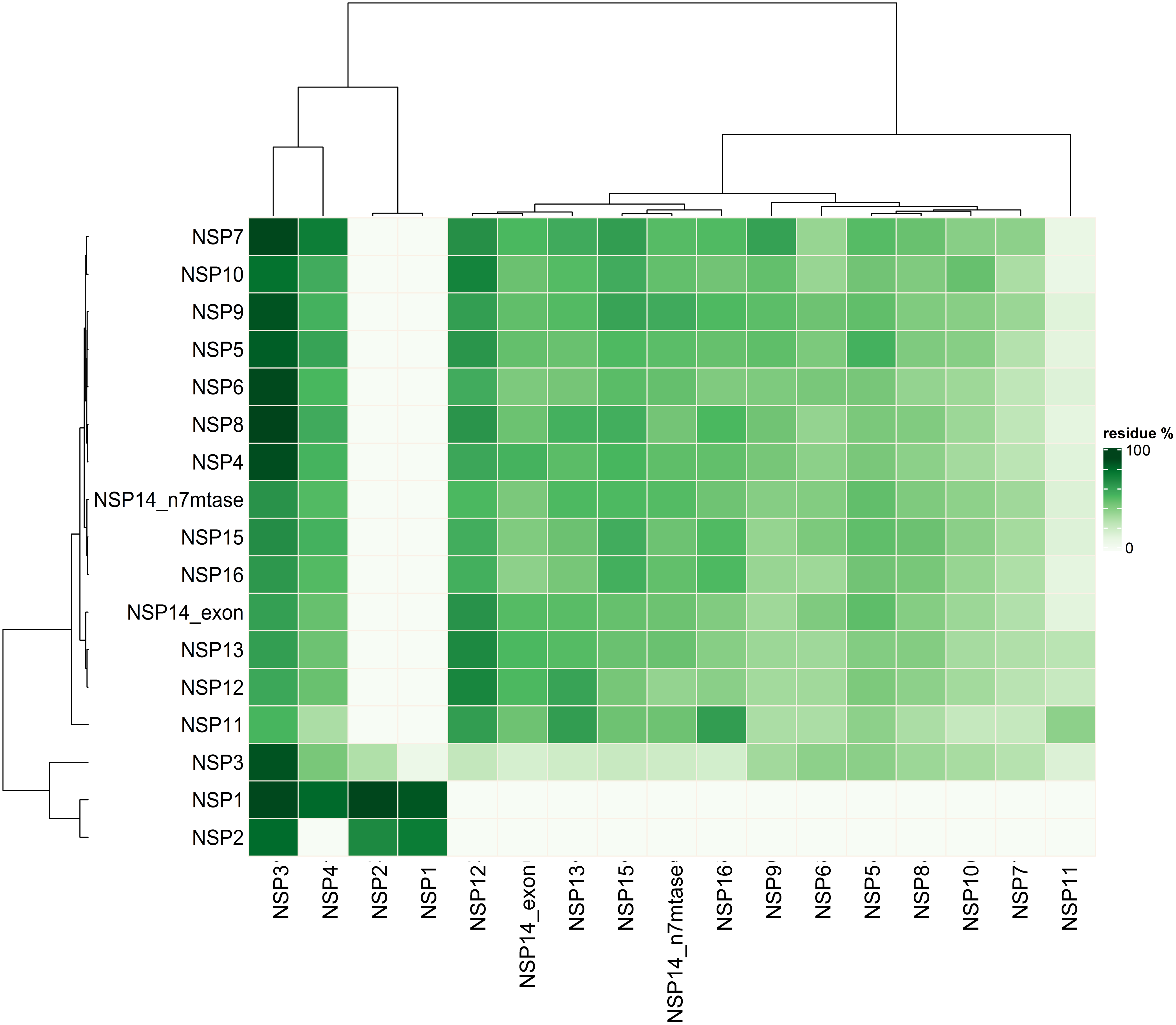

NSP7
NSP10
NSP9
NSP5
NSP6
NSP8
NSP4
NSP14_n7mtase
NSP15
NSP16
NSP14_exon
NSP13
NSP12
NSP11
NSP3
NSP1
NSP2
NSP3
NSP4
NSP2
NSP1
NSP12
NSP14_exon
NSP13
NSP15
NSP14_n7mtase
NSP16
NSP9
NSP6
NSP5
NSP8
NSP10
NSP7
NSP11
residue %
100
0
